# Systemic Calcitonin Gene-Related Peptide (CGRP) modifies auditory and vestibular end organ electrical potentials, and increases sensory hypersensitivities

**DOI:** 10.1101/2022.06.03.494764

**Authors:** Shafaqat M. Rahman, Stefanie Faucher, Raajan Jonnala, Joseph C. Holt, Choongheon Lee, Anne E. Luebke

## Abstract

Migraine is a severe and chronic neurological disorder that affects ∼18% of people worldwide, the majority being female (3:1). It is characterized by recurrent, debilitating headaches and heightened sensory sensitivities. People with migraine may develop vestibular migraine (VM), characterized by a heightened motion sensitivity and preponderance for spontaneous vertigo attacks and balance problems such as postural instability. Calcitonin gene-related peptide (CGRP) is implicated in migraine and is believed to act on brain meninges or in subcortical central nervous system (CNS) structures, and CGRP-based antagonists have shown efficacy for migraine treatment. CGRP also signals at efferent synapses of the cochlea and vestibular end organs, but it is unclear if exogenous CGRP can modulate inner ear function at the end organ level and cause heightened behavioral responses consistent with VM. We tested if intraperitoneally (IP) delivered CGRP to wildtype mice can modulate end organ potentials to sound [via auditory brainstem responses (ABRs)] and jerk stimuli [via vestibular sensory evoked potentials (VsEPs)]. We also assessed behavioral measures of phonophobia [acoustic startle reflex (ASR)] and static imbalance [postural sway-center of pressure (CoP)] after IP CGRP, and observed female mice exhibited heightened sensitivities to IP CGRP in all assays. Male mice showed similar auditory sensitivity and end organ effects to CGRP, but systemic CGRP did not modify male postural sway as it did in females. In conclusion, we show that intraperitoneally delivered CGRP affects ABRs and VsEPs, and elicits behaviors suggestive of auditory hypersensitivity and postural instability in mice related to the phonophobia and postural instability seen in VM patients.

**Significance Statement:** Calcitonin Gene-Related Peptide (CGRP) has been implicated in migraine, and CGRP-based therapeutics have shown efficacy in the treatment of migraine headaches. CGRP is also present in efferent synapses of the inner ear, so we questioned if increases in systemic CGRP can act directly on inner ear end organs and modify corresponding behavioral responses. In this study, we determined systemic CGRP changes cochlear and vestibular end organ potentials and produces migraine behaviors similar to phonophobia and postural control deficits. Peripheral changes in auditory and vestibular compound action potentials following systemic CGRP administration suggest CGRP can act directly on the inner ear, which provides one site of action for CGRP’s involvement in hypersensitivity to sound and movement in migraine.

## Introduction

Migraine is a debilitating chronic neurological disorder that is listed by the World Health Organization (WHO) as the most debilitating neurological disorder worldwide and is estimated to affect 18% of women and 6% of men (Goadsby, Lipton, & Ferrari, 2002; Lipton et al., 2007). About 30-50% of people with migraine have a vestibular component causing balance problems and dizziness (Kuritzky, Toglia, & Thomas, 1981; H. Neuhauser, Leopold, von Brevern, Arnold, & Lempert, 2001; H. K. Neuhauser, 2007, 2009) (Vukovic et al., 2007). Vestibular Migraine (VM) is a major cause of vertigo in dizziness clinics (Kayan & Hood, 1984) and patients with migraine, and especially VM, exhibit a heightened sense to sound and motion (Ashkenazi, Mushtaq, Yang, & Oshinsky, 2009; Lopez-Escamez et al., 2014). Despite migraine and VM’s prevalence, the pathophysiology remains uncertain (Lempert et al., 2022; Lempert et al., 2012).

Calcitonin gene-related peptide (CGRP) is intensively studied for its role in migraine’s sensory hypersensitivities (A. F. Russo, 2015). CGRP infusion evokes headache in migraine patients but not in non-migraineurs (Lassen et al., 2002), and CGRP levels are elevated during spontaneous migraine attacks (Goadsby & Edvinsson, 1993). Normalization of CGRP levels is associated with the relief of headache pain which is thought to act at either brain meninges or the peripheral nervous system (PNS) (Goadsby et al., 2002; Juhasz et al., 2005; Lassen et al., 2002)) . CGRP also signals at efferent synapses of the cochlea and vestibular end organs, so we questioned if elevated CGRP levels in migraine could also affect the inner ear (Dickerson, Bussey-Gaborski, Holt, Jordan, & Luebke, 2016; Luebke et al., 2014; Maison, Emeson, Adams, Luebke, & Liberman, 2003). We also questioned if increases in systemic CGRP can cause behaviors suggestive of sound hypersensitivity (phonophobia) or motion hypersensitivity, which are clinical symptoms observed in patients with migraine and VM.

In this study, we tested wildtype C57BL/6 mice to determine if there is a change in inner ear end organ potentials to sound (measured using auditory brainstem responses - ABRs) and jerk stimulus (measured using vestibular sensory evoked potentials - VsEPs) after an intraperitoneal (IP) CGRP injection. We also assessed corresponding behavioral measures of phonophobia (acoustic startle reflex – ASR) and imbalance (postural sway/center of pressure - CoP) after IP CGRP administration.

## Methods

### Animals

C57BL/6J mice were obtained from Jackson Laboratories (JAX #664) and were housed under a 12- hour day/night cycle under the care of the University Committee on Animal Resources (UCAR) at the University of Rochester. Mice were housed together in cages (5 mice per cage) with *ad libitum* access to food, water, and nesting and bedding materials. A total of 74 mice (37F/ 37M) were used for these studies, which were powered sufficiently to detect male/female differences. Before experimentation, mice were equilibrated in the testing room controlled for ambient temperatures between 22-23°C for at least 30 minutes and remained in this room until completion of experiments. IP injections occurred after the equilibration period. Mice used during behavioral experiments were returned to the vivarium, while mice used for ABR/VsEP experiments were euthanized after testing. Mice tested were between 2.3 - 6 months of age. This age range in mice correlates to 20-30 years in humans, which is within the age range that patients develop migraine symptoms (Lessem, 2018). All animal procedures were approved by the University of Rochester’s IACUC committee and performed in accordance with NIH standards. Moreover, the experimental protocols followed the guidelines for Animal Research Reporting In Vivo Experiments (ARRIVE) (McGrath & Lilley, 2015).

### VsEP and ABR Protocol

C57BL/6J mice were anesthetized with IP urethane/xylazine (1.2 g/kg/20 mg/kg). VsEP and ABR recording methods have been described in detail elsewhere (Lee, Holt, & Jones, 2017). Briefly, to record VsEP and ABR, subcutaneous electrodes were placed on the midline over the nuchal crest (G1), behind the right pinna (G2), and on the hip (ground). A noninvasive head clip was used to secure the mouse’s head to an electromechanical shaker. During VsEPs, the shaker delivered a transient (jerk) head translation in the naso-occipital axis (± X). To confirm the absence of auditory responses during VsEP testing, a binaural forward masker (intensity: 92 dB SPL; bandwidth: 50∼50 kHz) was presented with the use of a free-field speaker (FF1, TDT). For ABR recordings, the same electrodes were used to record ABRs, which were evoked by a single rectangular voltage pulse of 0.03 ms duration (corresponding to 33 kHz cut-off frequency). These click stimuli were presented via a free-field speaker, which was the same driver for acoustic masking during the VsEP recording. A noninvasive rectal thermometer or thermistor was used to monitor the animals’ core temperature at 38.0 ± 0.2°C. The standard signal averaging technique described in previous studies (Lee et al., 2017) was used to record both VsEP and ABR, as shown in **Fig. 1A**. Recordings of both ABRs and VsEPs were made in a soundproof booth. Both ABRs at a maximal intensity of 100 dB SPL, and VsEPs at a maximal intensity of +6 dB re: 1 gravity (g)/ milliseconds (ms), were recorded as a stable baseline before a single IP injection of either vehicle control (PBS) or 0.1 mg/kg CGRP was given. Control amplitudes and latencies are depicted as the averages of the 2-3 measurements taken during baseline before injection. Following single IP injections of either vehicle or CGRP, ABR and VsEP recordings were continuously measured where significant shifts in ABR and VsEP responses were first observed in 10 mins and reached a plateau within 15-20 mins of drug injection (post-CGRP; data not shown). Thus, amplitudes and latencies depicted as post-saline/post-CGRP are averages of the 2-3 measurements taken after 20-30 min post-injections.

**Fig. 1.**
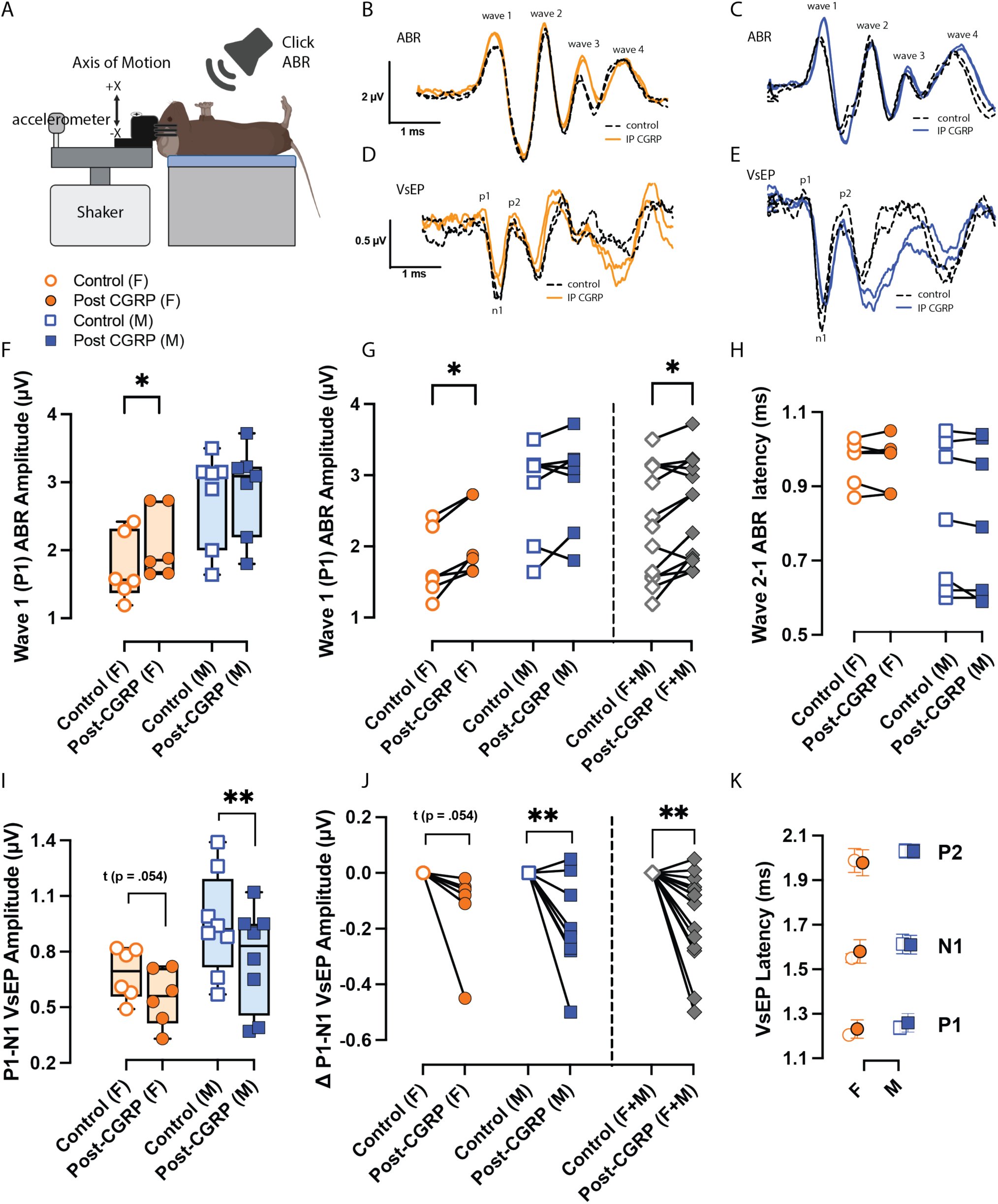
***A,*** Schematic illustrating mouse setup for VsEP/ABR recordings captured during baseline (control) and 20-30 minutes after intraperitoneal (IP) injections of either PBS as the vehicle or 0.1 mg/kg CGRP in female (F) and male (M) C57BL/6J mice**. *B-C***, Representative click-evoked ABR waveforms - in duplicate -were collected for ABR at a maximal intensity of 100 dB SPL, and waves were labeled (wave1 – wave 4). ***D-E,*** Likewise, representative VsEP waveforms were collected at +6 dB re: 1 g/ms having three, labeled response peaks (p1, n1, and p2). Orange and blue colors are used to distinguish female and male tracings post-CGRP, respectively. The following symbols and colors are used: female - orange, male - square, control - open, CGRP - closed. Sample sizes list - ABR (n = 6F/7M), VsEP (n = 6F/8M). ***G & J,*** Before/after plots indicate the change in amplitude post-CGRP compared to a control response normalized to zero. ***F-G***, A significant increase is seen in wave 1 ABR amplitudes post CGRP for female only, and when sexes are pooled. ***I-J*,** For VsEPs, a statistically significant decrease in P1-N1 VsEP amplitudes was seen in males only and when sexes were pooled, whereas a trend was seen in females only. IP CGRP had no significant effects on ***(H)*** ABR (wave 2-1) or ***(K)*** VsEP latencies. *p < 0.05*, p < 0.01**, p < 0.001***, p < 0.0001\*\*\*\**.

### Drug administration

Injections were performed intraperitoneally (IP) with a 30-gauge insulin syringe. Dulbecco PBS served as the vehicle control. IP CGRP was prepared at 0.1 mg/kg (rat ɑ-CGRP, Sigma). Injection volumes were calibrated so that each animal received 1 injection per testing day at ∼100 ul. For behavioral experiments, mice were tested ∼20-30 minutes after IP delivery of either vehicle or CGRP.

### Behavioral test of phonophobia: Acoustic startle reflex (ASR)

We used the acoustic startle response (ASR) to assess heightened sound sensitivity as a surrogate behavior for phonophobia in mice. Experiments were conducted within a sound-attenuating room (IAC, Bronx, NY) with Sonex foam lined walls. Mice were placed individually in a wire-mesh cage that is 5 cm wide, 7 cm long, and 4 cm high, and having free sound penetration. The testing cage was oriented so that the mouse faced a Tucker Davis ES2 electrostatic speaker (Masker Speaker) located 46 cm directly in front of the mouse’s head (60dB BBN). The startle speaker was a Yamaha JA4281B compression tweeter that was suspended 15 cm above the mouse (80-130 dB). The acoustic startle speaker and its supports, the pedestal, acrylic shelf, and the table on which the apparatus was placed were all covered with echo absorbing foam and carpeting. Startle eliciting stimuli (ES) were 15 ms broad-band noisebursts (5 ms linear-gating, 50 kHz bandwidth, 130 dB SPL) digitally generated using a Tucker-Davis Technology (TDT, Alachua, FL) RP2.1 Real-time Processor, attenuated using a TDT PA5, then amplified with an Adcom (East Brunswick, NJ) GFA-535 II amplifier and broadcast from the Startle Speaker above the mouse. Masker stimuli were digitally generated using a TDT-RP2.1 (100 kHz sample rate) and broadcast from the masker ES speaker. The force of the startle reflex was transduced by a custom accelerometer and the voltage output sampled at 1 kHz by the first RP2.1. The startle response amplitude was the RMS of this output in the 100 ms period after the delivery of the startle stimulus. The experiment was controlled from a PC using a custom Matlab (The Mathworks, Inc) front-end. Sound levels were measured with a ¼” microphone (Bruel & Kjær model 4135) connected to a measuring amplifier (Bruel & Kjær model 2610). Each testing session began with the mouse being placed within the testing cage in the startle chamber for a 2 minute acclimatization period, prior to delivery of stimuli. Each Acoustic startle (ASR) test sessions had 11 presentations of each condition, these being block randomized, and the inter-trial interval was randomized between 15 and 25 seconds. The responses from the first block were not analyzed to avoid potential large responses on the first few trials. There were 12 conditions in this session; startle stimuli were 80 to 130 dB SPL and delivered in silence or in a continuous background noise, which, when present, was on for the duration of the inter-trial interval and broadcast from the Masker Speaker. Mice were first tested 25 minutes after a vehicle (saline) injection, and then one week later, the same mice were retested 25 minutes after an IP injection of rat DCGRP (Sigma). The testing duration is 45 min/mouse, and ASR has excellent test/retest reliability (Allen & Luebke, 2017).

### Behavioral test of postural sway

Postural sway was examined in ambient light conditions (30 lux) using the center of pressure (CoP) test which acts as surrogate behavior for static imbalance in mice and has been evaluated in mouse models for neurodegenerative disease such as Parkinson’s and Alzheimer’s disease (Hutchinson et al., 2007). Mice were weighed and placed on a force plate designed to measure forces in the X, Y, Z axes and moments in the XY, YZ, and XZ directions. We used the AMTI Biomechanics Force platform (model HEX6x6) and AMTI automated acquisition software to obtain forces and moments and then MATLAB code was used to calculate CoP measures based on a confidence ellipse enclosing 95% of the CoP trajectory values recorded in a single trial. During a given test, mice were first allowed to freely maneuver on the force plate for 2-5 minutes to acclimate to the novel environment. An accessory plexiglass guard was placed around the force plate to prevent mice from moving off the test plate. After acclimation, ∼ 10 baseline trials were taken of mice with each trial indicating a CoP area (resolution per CoP measurement: 300 samples per second). These trials were taken when the mouse was not moving, and its four paws were touching the force platform. Mice were then subjected to a vestibular challenge (see section *<u>Vestibular Challenge: Orbital Rotation</u>* in methods) and were placed back onto the force platform. After a five-minute recovery period, an additional 10 trials were recorded of the mouse’s sway to determine post-VC effects on CoP as schematized in **Fig. 3**.

Mice were first tested after systemic delivery of IP vehicle (20 min after injection), with sway measured before and after the VC. Four days after vehicle testing, the same mice were injected with IP CGRP and were tested again 20 minutes after injection (timing correlated with end organ potential changes in ABRs and VsEPs). While systemic CGRP’s effect is believed to dissipate two to three hours post–injection, we retested after four days to avoid any residual dizziness due to VC or lingering CGRP effects. We also injected vehicle 2x to assess test/retest reliability and saw no differences.

Unreasonably large CoP values (> 15 cm^2^) suggested mice were either moving/grooming during the trial or that they were at the edge of the force plate, and those values were excluded based on a 10% ROUT analysis (GraphPad Prism 9.5). Even with values excluded, a minimum of eight different 95% ellipses were obtained per mouse in each testing condition.

A concern of using a plexiglass guard to keep mice on the force plate is that mice would potentially lean against the guard. In our experiences, mice did not lean on the plexiglass guard, as there was at least a 1 cm gap between the walls of the plexiglass guard and the force plate’s edges. This 1 cm gap was designed to ensure the guard and forced plate did not touch and generate friction that could be caught in sway measurements. Anecdotally, during IP vehicle experiments, we noticed instances where mice would gravitate to the edge of the force plate after being placed in the setup, and would leave their tail in the 1 cm gap between the guard and force plate. In these cases, the force plate was reset and mice were re-positioned to the center. If a mouse repeatedly moved to the edge of the force plate during testing, then this mouse was not further tested on that day. This mouse would be re- tested at a later part of the day, or at a later day in the week. We did not have this issue with IP CGRP testing.

### Vestibular Challenge: Orbital Rotation

In a previous paper from our group, we challenged dynamic balance behaviors in mice, and used off-vertical axis rotation (OVAR) as the vestibular challenge, but noticed this stimulus caused significant anxiety in mice (Rahman, Hauser, Faucher, Fine, & Luebke, 2024). When we evaluated OVAR’s effects on postural sway using the CoP assay, we observed mice to have very small ellipse areas post-OVAR and freezing behavior (unpublished), further suggesting significant anxiety. Thus, we decided to determine a less stress-inducing vestibular provocation for postural sway testing. In this study, mice were subjected to a vestibular challenge in which they were rotated at 125 rpm (orbital displacement = 2 cm) at no tilt for 5 minutes. The mouse’s body and head are not fixed to allow for free head movements. Anecdotally, we did not observe anxiety-related behaviors in mice after this challenge.

Vestibular provocation has been explored by many groups to elicit motion sensitivity and motion sickness in rodent models, and we are not the first to do these types of studies. However, prior studies use very intense rotational stimuli to induce motion sickness in mice (Idoux, Tagliabue, & Beraneck, 2018; Li, Zhang, Zheng, & Huang, 2008; Yu, Cai, Liu, Chu, & Su, 2007) which we decided were not suitable for this study.

### Statistical Analyses

All statistical analyses were conducted in GraphPad Prism 9.5. Analyses were conducted separately in females and males. Repeated measure ANOVA (RM-ANOVA) and paired t-tests were the primary statistical tools. Tukey post hoc analysis was used to assess treatment differences in females or males, and Bonferroni post hoc analysis was used to assess treatment differences between sexes. Significance was set at *p* < 0.05 for all analyses.

## Results

### Effect of IP CGRP on ABR

Representative traces are shown for ABR recordings in female and male C57BL/6J mice **(Fig. 1B, C)**. Female control ABR wave 1 (P1) amplitudes (*n* = 6) were computed to be 1.74 ± 0.20 µV and increased to 2.08 ± 0.21 µV after IP CGRP (one-tailed paired t-test; *t, df* = 3.94, 5; *p* = 0.01). In males, control ABR wave 1 amplitudes (*n* = 7) were computed to be 2.78 ± 0.26 µV and changed to 2.89 ± 0.25 µV after IP CGRP (one-tailed paired t-test; *t, df* = 1.02, 6; not significant) **(Fig. 1F)**. Wave 1 ABR amplitudes were normalized so that post-CGRP measurements highlight the change in amplitude from their respective control measurements (Δ wave1 amplitude) **(Fig. 1G)**. When sexes are pooled (*n* = 13), a significant Δ increase of 0.22 ± 0.08 µV was observed in wave 1 amplitudes due to IP CGRP (one-tailed paired t-test; *t, df* = 2.89,12; p = 0.01). In parallel, a different group of mice were tested to examine the effects of the vehicle control, IP PBS, on the ABR response (**Supplementary Fig. 1A, B**). IP vehicle induced negligible changes in ABR wave 1 amplitude, as changes were regarded not significant in females (n = 4) or males (n = 4). Wave 2-1 ABR latencies were calculated as the difference between P2 and P1 latencies for control and post-CGRP; however, no differences were observed in latencies **(Fig. 1H).** IP vehicle also had no effects on Wave 2-1 ABR latencies (**Supplementary Fig. 1C)**.

### Effect of IP CGRP on VsEPs

Like ABR tests, representative traces are shown for VsEP recordings in females and males (**Fig. 1D, E).** Female P1-N1 VsEP control amplitudes (*n* = 6) were computed to be 0.68 ± 0.06 µV and were 0.55 ± 0.06 µV after IP CGRP and deemed not statistically significant (one-tailed paired t-test; *t, df* = 1.96, 5; *t* (*p* = 0.054)) **(Fig. 1I)**. Likewise, male P1-N1 VsEP control amplitudes (*n* = 8) were computed to be 0.95 ± 0.10 µV but were reduced significantly to 0.76 ± 0.10 µV after IP CGRP (one-tailed paired t-test; *t, df* = 3.00, 7; *p* = 0.009). P1-N1 amplitudes were normalized so that post-CGRP measurements highlight the change in amplitude from their respective control (Δ P1-N1 amplitude), and both sexes were additionally pooled (**Fig. 1J**). When pooling all animals (*n* = 14), a significant Δ reduction of -0.16 ± 0.04 uV in P1-N1 amplitude is observed due to IP CGRP (one-tailed paired t-test; *t, df* = 3.65, 13; *p* = 0.002). These changes translate to an 18.9% reduction in VsEP P1-N1 amplitude in females and a 19.4% reduction in males. In contrast, IP vehicle did not lead to statistically significant changes in VsEP P1-N1 amplitude for females (n = 5) or males (n = 4) (**Supplementary Fig. 1D, E**). P1, N1, and P2 latencies were also assessed for VsEPs, and these latencies are depicted as mean ± SEM for females and males during control and post-CGRP tests **(Fig. 1K),** with no significant difference due to IP CGRP. IP PBS had no significant effects on P1, N1, or P2 latencies (**Supplementary Fig. 1F**).

### CGRP’s effect on sound sensitivity

Our sound sensitivity measure in the mouse model makes use of the acoustic startle response (ASR) in quiet and in background noise. Mice were assessed for the effects of IP CGRP as compared to IP vehicle (serving as baseline). Mice were first tested for their ASR in quiet and in background noise ∼30 min after a vehicle (saline) injection, and then one week later, the same mice were re-tested 30 min after an IP injection of CGRP. The ASR experimental setup is illustrated in **Fig. 2 A** (quiet) and **Fig. 2D** (background noise 60 dB). In both IP vehicle and IP CGRP injected animals, an increase in ASR was detected with increasing loudness of the startle stimulus (ES). All animals tested exhibited robust startle responses (*n* = 6F/ 6M) to vehicle injections. When ASR was tested after IP CGRP injection and compared to IP vehicle, all animals exhibited reduced ASR thresholds in both quiet and noise. IP CGRP significantly increased sound sensitivity to the startle stimulus in quiet (**Fig. 2 B, C**) and in background noise (**Fig. 2 E, F**) in both female and male mice at all sound levels (****= *p* < 0.0001 quiet, ***= *p* < 0.001 in noise).

**Fig. 2.**
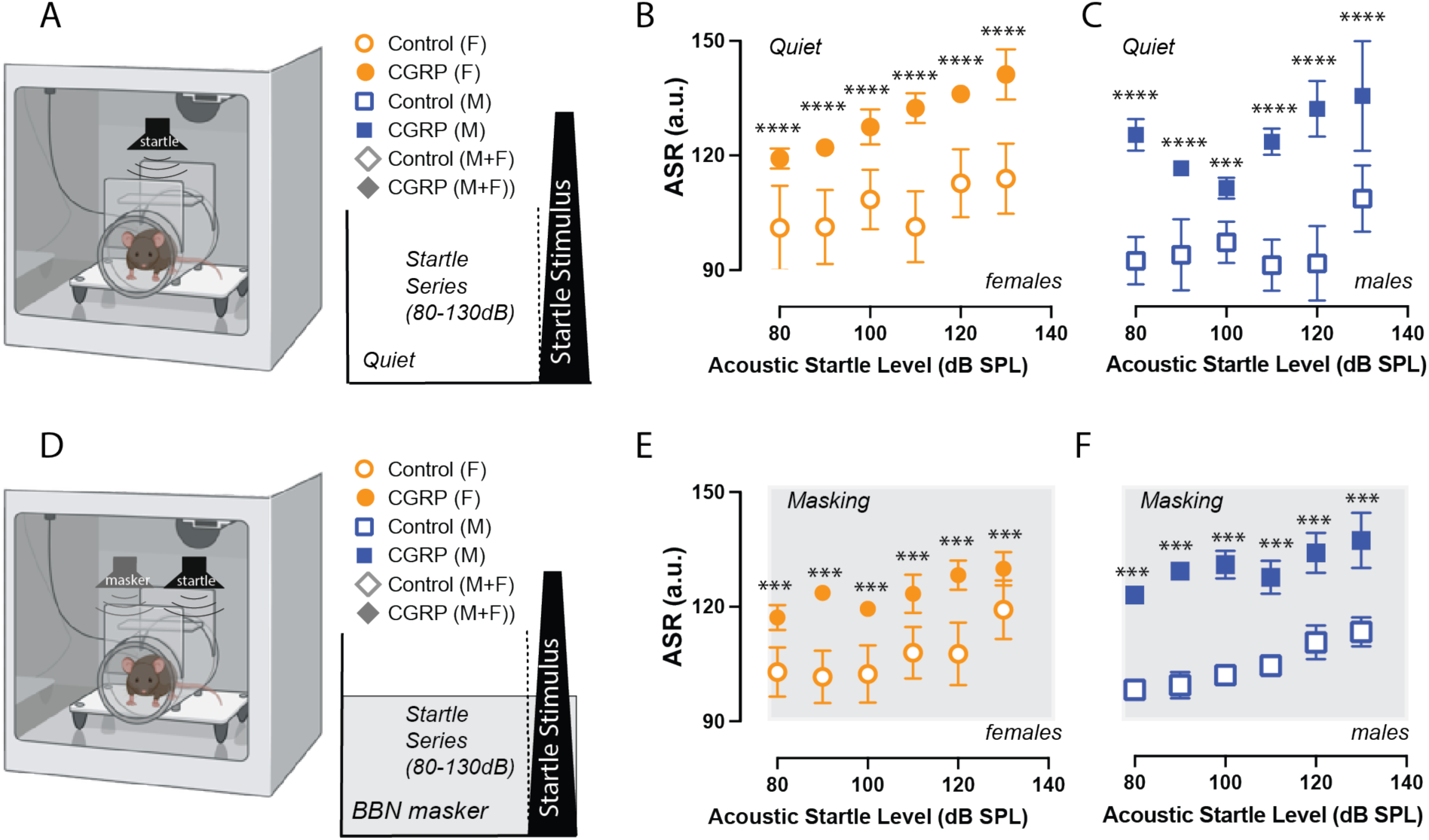
Systemic CGRP increases Acoustic Startle Responses (ASR) in both quiet and background noise, suggesting increased sound sensitivity. *A,* Schematic of startle stimulus presented in otherwise quiet background conditions. *B-C*, in female mice (*B*, n = 6), and male mice (*C*, n=6), there were significant differences between vehicle and IP CGRP ASR measures in quiet conditions (p < 0.0001). All mice were more sensitive to the same sound level of ASR stimulus after being injected with CGRP. *D,* Schematic of startle stimulus embedded in continuous 60dB SPL broadband background noise. *E-F*, ASR embedded in noise was also more sensitive to same sound level after CGRP injection as shown for female mice (*E*, n = 6), male mice (*F*, n = 6), (p < 0.001). All mice were more sensitive to the same level of ASR stimulus whether delivered in quiet or in BBN after being injected with CGRP. Moreover, all mice showed ASR increases with sound level from 80 to 130dB SPL with near maximal startle is elicited by 130 dB SPL stimulus levels. Error bars are SEMs. *P < 0.05*, p < 0.01**, p < 0.001***, p < 0.0001\*\*\*\**.

**Fig. 3.**
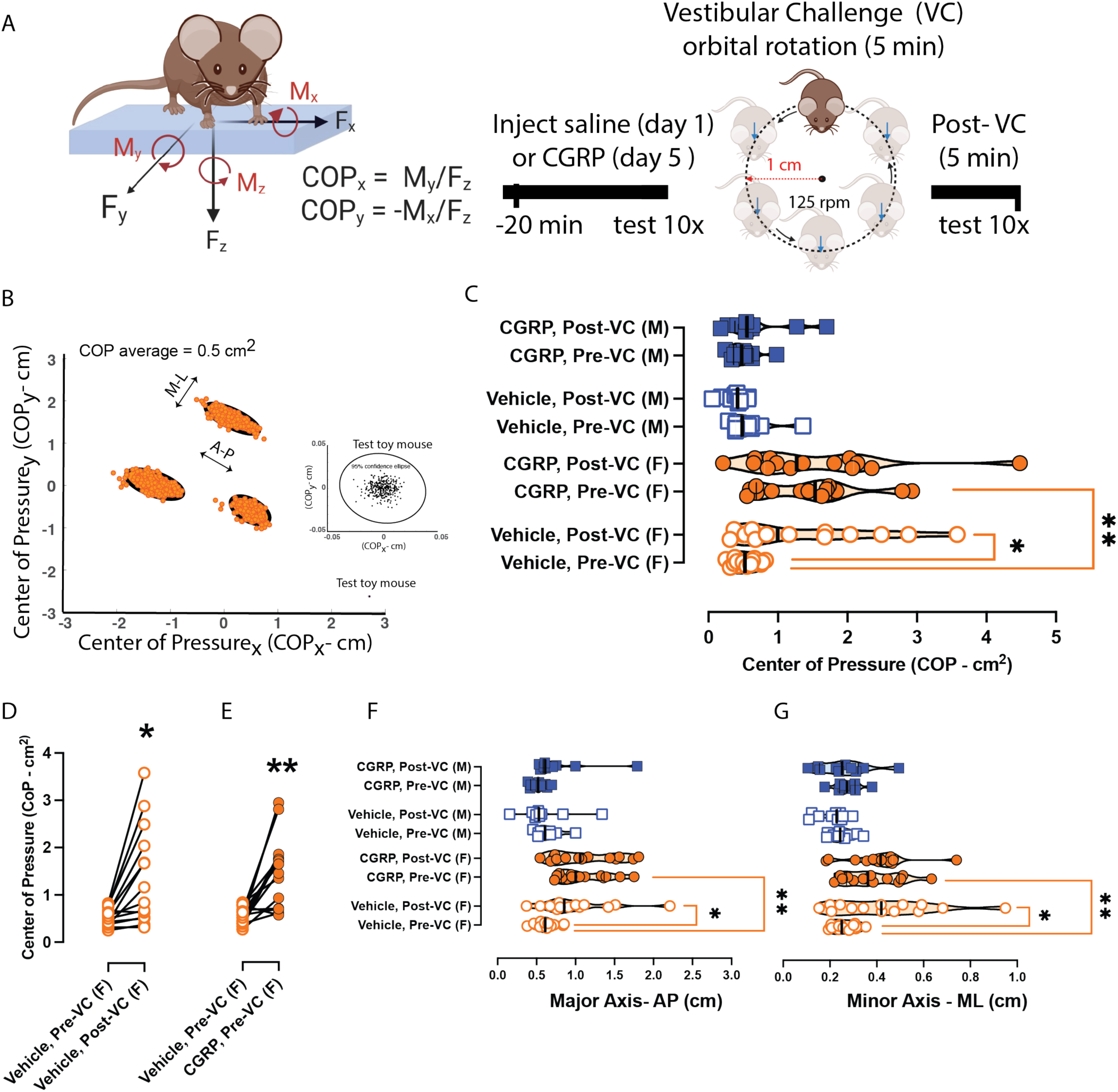
CGRP and Vestibular Challenge increase postural sway in female mice. Symbols and colors are depicted in a similar scheme as previous figures. ***A,*** Timeline involved mice being assessed for effects of IP PBS or IP CGRP on day 1 or day 5, respectively. A 5-minute orbital rotation (125 rpm for 5 minutes, orbital radius = 1 cm) was used as the vestibular challenge, and mice were repeatedly tested (n = 14F/10M). ***B,*** Three distinct 95% ellipse areas during pre-VC testing were computed to be 0.50 ± 0.06 cm^2^ for an example female mouse. In the inset of **B**, a toy mouse was measured to be 0.02 ± 0.0005 cm^2^ for comparison with live animals. ***C,*** IP CGRP increased female postural sway but had little effect in males. **D**, Females exhibit greater sway after vestibular challenge and after vehicle injection. ***E,*** Before-after plot is shown to depict changes in sway in repeatedly tested animals. ***F &G,*** Prior to the VC, IP CGRP induced longer AP and ML ellipse lengths in females than was observed during their IP vehicle testing. In addition in females, the vestibular challenge was observed to induce longer AP and ML lengths during IP vehicle testing but not after IP CGRP, which is consistent with CoP area changes. *p < 0.05*, p < 0.01**, p < 0.001***, p < 0.0001\*\*\*\**.

### CGRP’s and VC’s effects on postural sway

Mice were assessed for postural sway by determining center of pressure (CoP) ellipse area. An experimental timeline is shown in **Fig. 3A**. Twenty minutes after delivery of IP vehicle and without a vestibular challenge (VC), group averages of 95% ellipse areas were computed. An example of three distinct 95% confident ellipse areas (3 trials) for one female mouse tested on postural sway about twenty minutes after IP vehicle administration is shown if Fig. **3B**, to give a visual of what each trial looks in respect to the location of the mouse on the platform. There is little variability in each trial regardless of where the mouse is on the platform. The inset in **Fig. 3B**, shows the ellipse when we placed a toy mouse at the center of the platform (0 cm/0 cm) to act as a still, motionless weight when compared to a live mouse, and this still weight generated stable and tiny 95% confident ellipse areas. The toy mouse’s average 95% CoP ellipse area was computed to be 0.02 ± 0.0005 cm^2^. When we moved this toy mouse to different areas on the platform, we generated near identical outputs. A cohort of mice (*n* = 10F/ 10M) was assessed for a test-retest of 95% ellipse areas after IP vehicle treatment, and testing occurred four days apart. A 2-way repeated measure ANOVA with Bonferroni post-hoc analysis indicated no significant differences in either sex during the test-retest for females (mean difference = 0.10 ± 0.3 cm^2^) or for males (mean difference = -0.14 + 0.3 cm^2^) (p > 0.99 for either sex). In either sex, 2-way ANOVAs with Tukey post hoc analyses were used to assess the factors of CGRP and VC on 95% CoP ellipse areas in mice (n = 14F/ 10M) **(Fig. 3C).** Differences were seen in females due to IP CGRP (*F* (1.00, 13.00) = 4.77, *p* = 0.05) and the VC (*F* (1.00, 13.00) = 4.49, *p* = 0.05). In females after IP vehicle, the VC induced higher CoP ellipse compared to pre-VC trials (*p* = 0.02) as shown in **Fig. 3D**. When comparing pre-VC trials, IP CGRP induced increases in female CoP areas by 0.94 ± 0.23 cm^2^ compared to IP vehicle (*p* = 0.003) as seen in **Fig. 3E**. Females also had 0.96 ± 0.15 cm^2^ higher CoP areas than males based on a separate 2-way ANOVA assessing sex differences in pre-VC, CGRP and post-VC, CGRP results (*p* = 0.001).

The major and minor axes of 95% ellipse areas were also analyzed with 2-way ANOVAs across the factors CGRP and VC (**Fig. 3F** and **3G**). The major and minor axes lengths serve as correlates for anteroposterior (AP) and mediolateral (ML) sway magnitudes. A 2-way RM ANOVA on major axes indicated significant effects in females due to IP CGRP (*F* (1.00, 13.00) = 5.59; *p* = 0.03) and VC (*F* (1.00, 13.00) = 5.41; *p* = 0.04) whereas these effects were not observed in males. In comparison, ANOVAs that analyzed minor axes indicated a VC effect in only females (*F* (1.00, 13.00) = 7.46; *p* = 0.02). Tukey post-hoc indicated IP CGRP induced larger AP ellipse lengths by 0.49 + 0.10 cm in females compared to their IP vehicle trials (*p* = 0.001). Also, in IP vehicle-treated females, there was an increase in AP ellipse lengths during the post-VC test versus pre-VC (*p* = 0.02). IP CGRP-treated females exhibited greater AP lengths compared to males during pre-VC testing by 0.57 + 0.08 cm (*p* < 0.001). ML lengths increased after the VC in IP vehicle-treated females (*p* = 0.02), and IP CGRP-treated females had higher ML ellipse lengths compared to their IP vehicle response (*p* = 0.007). No other significant changes were observed in AP and ML sway.

## Discussion

Using electrophysiological techniques (ABR/VsEP), we observed that intraperitoneally delivered CGRP modulates end organ function in the inner ear with the maximum effect occurring ∼20- 30 minutes after injection. Exogenous CGRP increases wave 1 amplitudes of the ABR but reduces VsEP P1-N1 amplitudes, suggesting at least one action of systemic CGRP is at efferent targets of the auditory and vestibular periphery. Wave 1 is widely understood to arise respectively from the distal and proximal auditory nerve (Britt & Rossi, 2005; Kamerer, Neely, & Rasetshwane, 2020), whereas P1-N1 VsEP arise from peripheral vestibular nerve activity (T. A. Jones et al., 2011; Konrad-Martin et al., 2012).

These structures begin in the inner ear and travel to the brainstem and project to the cochlear and vestibular nuclei. Yet, as suggested by other studies, CGRP and its receptor are widely distributed across the cerebellum, thalamus, and hypothalamus, and CGRP’s effects in vestibular symptoms of migraine may be attributed to changes in the CNS such as the vestibulo-thalamo-cortical network (Espinosa-Sanchez & Lopez-Escamez, 2015; Lopez-Escamez et al., 2014; Warfvinge & Edvinsson, 2019).

Furthermore, we observed that IP CGRP - during this 20-30 minute timeframe - increased sound sensitivity in male and female C57BL/6J mice. Both the increases in ABR wave 1 amplitudes and the increased ASR responses are consistent with CGRP’s effect on compound action potentials (CAPs) observed in intra-cochlear infusion of CGRP(Le Prell, Hughes, Dolan, & Bledsoe, 2021). In addition, the increased sound sensitivity and increased wave 1 ABR amplitude with IP CGRP injection complements findings of decreased sound sensitivity and reduced wave 1 amplitude in DCGRP–null mice (Allen & Luebke, 2017; Dickerson et al., 2016; Maison et al., 2003). Thus, the auditory nerve may be hypersensitized to CGRP’s effects during a migraine.

Several studies have clearly demonstrated that P1-N1 VsEP arises from the peripheral vestibular nerve innervating the otolithic organs, and these studies involved far-field recording. Inner ear labyrinthectomies eliminated vestibular responses, whereas cochlear ablation failed to eliminate VsEP responses in birds and mammals (S. M. Jones, Jones, & Shukla, 1997; T. A. Jones, 1992; S. M. Jones, Erway, Bergstrom, Schimenti, & Jones, 1999; T. A. Jones & Jones, 1999). While acoustic masking eliminates ABRs, VsEPs persist under the same conditions. In a pertinent study in the chicken, brainstem sectioning of the second-order neuron eliminated most major P2 and later peaks, leaving P1 and N1 peaks intact, and suggesting P2 peak reflects activity of the first brainstem relay (Nazareth & Jones, 1998). In brief summary, this study provides clear evidence that VsEP peaks later than N1 are dependent on central relays, whereas P1 and N1 reflect peripheral activity. Additionally, a genetically engineered mouse model missing otoconia in the otolith organs also failed to show VsEPs, while all mice demonstrated robust ABRs (S. M. Jones et al., 1999). The same genetically mutant animals showed normal primary vestibular afferents (Hoffman, Ross, Varelas, Jones, & Jones, 2006; T. A. Jones, Jones, & Hoffman, 2008). Pharmacological applications of neurotoxins eliminate VsEPs through perilymphatic perfusion and round window application in the inner ear (Irons-Brown, Jones, & Jones, 2003; T. A. Jones, 1992).

Intraperitoneal (IP) injection of CGRP has been extensively studied in preclinical rodent models of migraine where it is believed to affect the peripheral nervous system (PNS). For instance, IP CGRP resulted in light-aversive behaviors in WT mice (Mason et al., 2017). Additionally, peripherally administered CGRP can produce spontaneous pain in mice as assessed by the grimace assay (Rea et al., 2018) . Our ABR and VsEP findings show that IP CGRP affects inner ear signaling. While the exact mechanism of how CGRP alters inner ear function is currently uncertain, it may represent the direct effects of CGRP on receptors in the inner ear following the perilymphatic entry of CGRP following systemic administration. While the entry of substances into the inner ear is not strictly regulated by the rules governing entry of that same substance into the CNS, the molecular weight (i.e., size) of that substance, its solubility measures, and whether that substance is charged can influence its access to inner ear fluids, particularly following systemic administration (Lee et al., 2021; Salt and Hirose, 2018). IP administration of AMPA receptor blockers (IEM1460 and IEM1925) has been shown to reach and modulate the inner ear despite being positively charged compounds, as evidenced by the presence of these blockers in perilymph samples and their effect in reducing CAP amplitudes (Walia et al., 2021). Furthermore, several charged cholinergic compounds can enter the inner ear despite their inability to cross the CNS BBB (Lee et al., 2021). Antisense nucleotides have also been shown to enter the perilymph following systemic administration (Lentz et al., 2013).

We also considered that another explanation for CGRP’s effects may be due to CGRP-mediated changes in inner ear blood flow (Hillerdal & Andersson, 1991; Quirk, Seidman, Laurikainen, Nuttall, & Miller, 1994), yet when CGRP was eliminated in CGRP-null mice, distortion product otoacoustic emissions (DPOAEs) which are very dependent on blood flow, were not affected (Dickerson et al., 2016). In fact, peripheral CGRP’s effect on light aversion was also independent of any vasodilatory effects as CGRP-induced light aversion was observed with normalized blood pressure (Mason et al., 2020).

In comparison to IP CGRP neuropeptide modulating inner ear activity, we did not observe significant changes at wave 1/P1-N1 in a separate group of mice tested with IP PBS as the vehicle control. We assessed IP PBS’s effects at the same time frame as we studied IP CGRP and observed no significant changes from baseline ABR/VsEP responses. Notably, our finding of IP CGRP reducing VsEP P1-N1 amplitude is a novel contribution to our study, while the trend of IP CGRP enhancing ABR is consistent with another study that demonstrated a 20-30% increase in CAP amplitude following intracochlear CGRP infusion (Le Prell et al., 2021). In our analysis of electrophysiological experiments where we pooled male and female responses together, we observed that IP PBS decreased mean ABR wave 1 by 4.0% and mean VsEP P1-N1 by 1.0% compared to their baseline control measurements. In contrast, IP CGRP resulted in a 9.3% increase in mean ABR wave 1 and a 19.4% decrease in mean VsEP P1-N1 from their baseline control measurements. Notably, IP CGRP had a profound effect on females by increasing mean ABR wave 1 amplitudes by 19% from their baseline tests. Several previous studies have shown normal saline, PBS and other vehicle solutions (DMSO, Kolliphor, and alcohol-based solutions) do not produce significant changes in VsEP/ABR responses over a duration of up to 2 hours (Lee et al., 2017; Lee & Jones, 2018, 2021) and our current IP PBS results strongly align with these earlier findings, thus reinforcing the reliability of our findings and providing confidence that IP CGRP is modulating inner ear afferent activity.

A limitation of our study is that we did not directly measure CGRP levels in the inner ear and could not directly attribute its role to causing ABR/VsEP changes, or to a potential downstream effect of increased CGRP levels in circulation. A future study for our group is to evaluate electrophysiology after delivering CGRP locally to the inner ear, using intracochlear delivery through the round window (Plontke, Hartsock, Gill, & Salt, 2016) or intracanal injections (Lee et al., 2021).

Besides our study which evaluates IP CGRP’s effects on the inner ear, a few studies have gauged CGRP’s role in vestibular dysfunction by evaluating CGRP expression in the brainstem’s vestibular nucleus. Intracerebral-ventricular (ICV) injection of nitroglycerin (NTG) caused impaired balance beam walking in rats. Immunohistochemical staining indicated heightened CGRP expression in the vestibular nucleus of the NTG rat model than in littermate controls. Moreover, intracerebroventricular administration of a CGRP-receptor antagonist (olcegepant-BIBN4096BS) into the left lateral ventricle was shown to be effective in alleviating impaired balance beam walking (Tian et al., 2022; Zhang et al., 2020). CGRP inhibition via olcegepant improved balance beam activity when delivered into the central nervous system, and so a future study for our group will be to evaluate CGRP inhibition using olcegepant delivered peripherally, either using IP injections as in this study, or more locally using intra- cochlear/round window injections.

Our postural sway testing showed a robust sex difference, as female mice were much more affected than their male counterparts to IP CGRP. In all mice IP CGRP injection was compared to each mouse’s baseline IP vehicle, with some female mice showing greater effects than others, but all showed increased postural sway to IP CGRP. It is unclear why we did not find significant changes in males during postural sway testing, yet female-specific sexual dimorphism has been observed in other preclinical assays assessing CGRP’s effects in migraine, but we are the first to show such differences in a preclinical postural sway assay (Paige et al., 2022; Recober et al., 2009).

Another limitation in this study is that we did not control for estrus cycle effects. However, the likelihood of these effects to be observed in the vestibular performance of humans is not currently supported by strong evidence, and data in mouse models is limited. Menstrual cycle timing has no effect on optokinetic function or anteroposterior sway in young women (Darlington, Ross, King, & Smith, 2001), and young women evaluated for changes in vestibulo-ocular reflex (VOR) function across their menstrual cycle phases showed no evidence of gain changes or asymmetry (Sinha, Mohamad, & Penwal, 2021). Anxiety is a risk factor for dizziness symptoms (Omara, Basiouny, Shabrawy, & Shafei, 2022), and the effect of estrous cycle on anxiety has been a topic of study in rodents, but results are mixed. Some studies suggest differences in mouse anxiety between proestrus, estrus, and diestrus phases (Frye, Petralia, & Rhodes, 2000; Gangitano, Salas, Teng, Perez, & De Biasi, 2009), whereas other studies showed no effect (Chari, Griswold, Andrews, & Fagiolini, 2020). We did test for rest- retest reliability on our SWAY and ASR assays and did not find any differences in mice responses when tested 1 week to 2 weeks apart. Moreover, e we did not find studies in mice evaluating auditory and vestibular electrophysiology or behaviors across estrus cycle, so future experiments addressing this concern are warranted.

Systemic administration of small molecule blockers of the CGRP receptor called ‘gepants’ have been shown to enter the peripheral nervous system like CGRP (Kim et al., 2022). The current evidence in favor of anti-CGRP monoclonal antibodies (mAbs) for treating VM is compelling. A prospective observational cohort study over 18 months observed significant reduction in vertigo and headache frequency in 50 VM patients treated with various anti-CGRP monoclonal antibodies (C. V. Russo et al., 2023), suggesting CGRP inhibition may an effective treatment option for VM patients who do not find relief from existing therapies. Future studies will make use of these preclinical electrophysiological and behavioral assays of auditory and vestibular components of CGRP-induced migraine symptoms to determine if CGRP receptor antagonism can relieve these auditory and vestibular symptoms and reduce ABR and VsEP responses.

## Conflict of interest statement

The authors declare no competing financial interests.

### Acknowledgments

This work was fully supported by NIH R01DC017261 (AEL).

---

**Supplementary Fig. 1.**
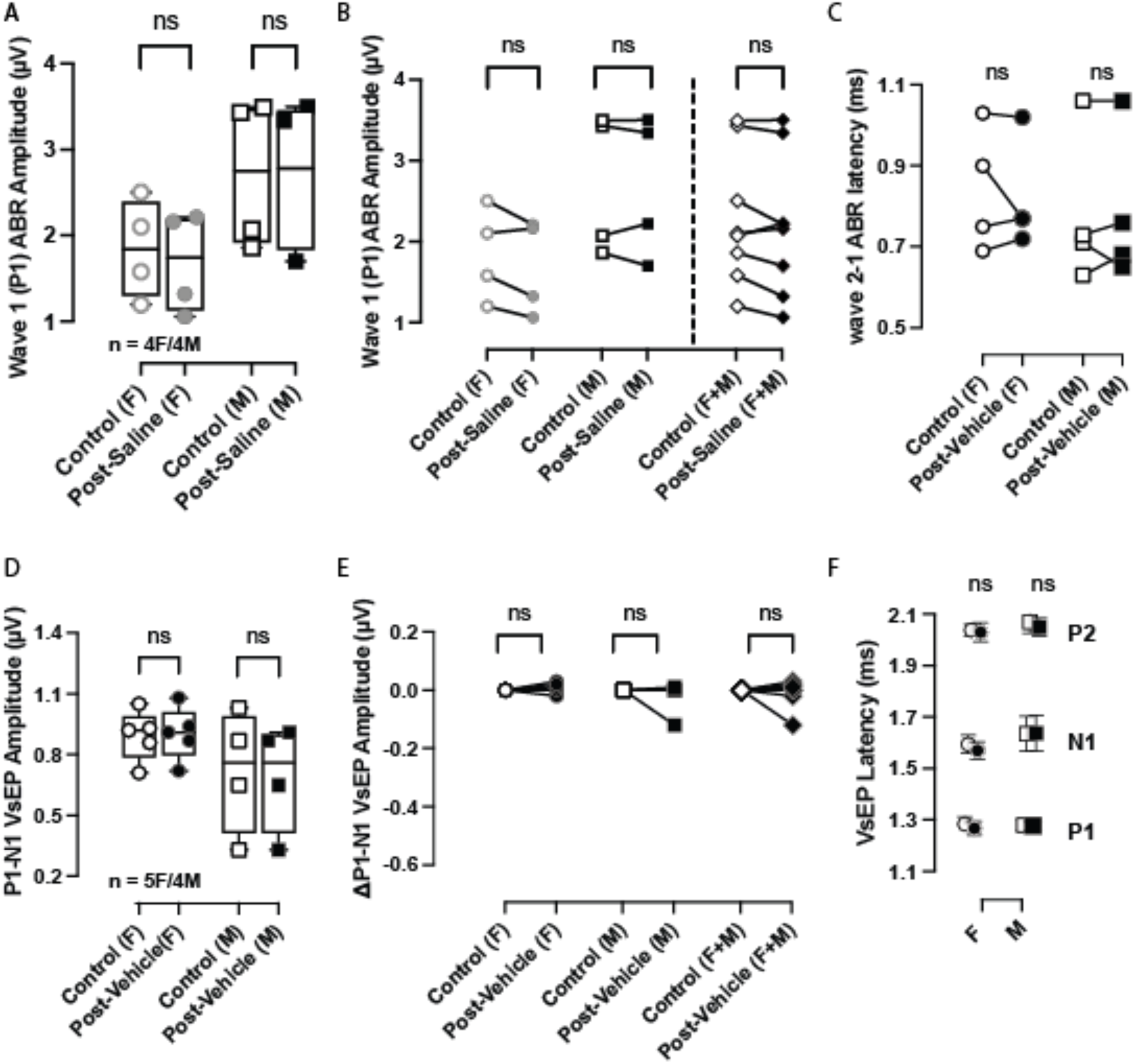
IP PBS (vehicle) was assessed for ABR (n = 4F/4M) and VsEP (n = 5M/4F) amplitude and latency responses. Females = circles, males = squares, and females + males = diamonds. Open circles indicate control measurements taken before the vehicle was administered, and post-vehicle measurements were taken 20 to 30 minutes after IP administration. One-tailed t-tests were used to assess differences in amplitudes between control (baseline) and post-vehicle for ABR (***A***) and VsEP (***D***). ***B&E***, The Δ change from control to post-vehicle for wave 1 ABR and P1-N1 amplitudes are represented as before-after plots. ***C&F***, Two-way RM ANOVAs were used to compare latency differences for ABR and VsEPs, but no differences were observed. *p < 0.05*, p < 0.01**, p < 0.001***, p < 0.0001\*\*\*\**.

